# FlcE latches onto the FliL-stator complex to turbocharge flagellar motility in *Borrelia burgdorferi*

**DOI:** 10.64898/2026.05.29.728556

**Authors:** Md Khalesur Rahman, Wangbiao Guo, Jack M. Botting, Michael Y. Galperin, Jun Liu, Md A. Motaleb

## Abstract

Periplasmic flagella are essential for the distinctive morphology and motility of the Lyme disease spirochete *Borrelia burgdorferi*, and motility plays a critical role in its pathogenic lifestyle. These flagella are powered by specialized motors that contain a large spirochete-specific multiprotein collar complex, yet the molecular architecture and mechanisms underlying high-torque motility remain poorly understood. Here, we identify the tetratricopeptide repeat (TPR)-containing protein BB0298 as a previously unrecognized flagellar collar component and rename it FlcE. Loss of *flcE* results in altered morphology and nearly abolishes the spirochete’s motility. Using cryo-electron tomography and biochemical analyses, we show that FlcE occupies a unique position within the collar where it surrounds the FliL-stator complex by binding to FliL and the collar protein FlcA. These findings support a model in which FlcE functions as a molecular “latch” that secures stator assemblies to sustain efficient torque generation. Notably, *flcE* is conserved across most motile members of the Spirochaetales and is located within the division and cell-wall (*dcw*) gene cluster. More broadly, the conservation of TPR-containing structural proteins across diverse bacterial flagellar systems suggests a general architectural principle whereby dedicated scaffolds reinforce stator complexes to maximize motor performance under high mechanical loads.

**Importance:** Spirochetal motility requires the stable engagement of torque-generating stator units, yet the mechanisms that secure these complexes have remained unknown. Here, we identify FlcE as a previously unrecognized collar protein that functions as a molecular latch by restraining the FliL-stator complex. Loss of FlcE destabilizes FliL-stator assemblies and severely impairs motility, demonstrating that peripheral scaffolding is essential for sustaining high torque output in the Lyme disease spirochete *Borrelia burgdorferi*. The conservation of tetratricopeptide repeat domains in FlcE and other bacterial flagellar proteins further suggests a broader mechanism in which specialized scaffolds anchor and stabilize stator complexes, enabling high torque generation. Because motility is a central virulence determinant in many pathogenic bacteria—including *B. burgdorferi* during transmission from the tick to the mammalian host and subsequent infection—defining the mechanistic role of FlcE provides important insight into the molecular basis of motility-driven pathogenesis and suggests new strategies for disrupting bacterial infection.

## Introduction

Flagellar motility is crucial for the survival and virulence of many pathogenic bacteria, including spirochetes (1–7). The flagellum consists of a motor, hook, and filament (8). The motor is a cell-envelope-spanning nanomachine composed primarily of a rotor ring and multiple stator complexes that convert ion-motive force into torque (9–11). Torque generated by the motor is transmitted through the hook to the filament, driving rotation and motility (12). Although the overall organization of bacterial flagella is broadly conserved, spirochetes possess a highly specialized periplasmic flagellum, which is located between the outer membrane and inner membrane (13–18). Each periplasmic flagellum is anchored near one cell pole and extends toward the opposite pole while wrapping around the cell body. Although the number and arrangement of periplasmic flagella vary among spirochete species, spirochetal motility is uniquely characterized by the rotation of the entire cell body, driven by the periplasmic flagella. (13).

The Lyme disease spirochete *Borrelia burgdorferi* has emerged as an ideal model system for understanding the distinctive structure and function of periplasmic flagella. At both cell poles, 7-11 periplasmic flagella form flat ribbons along the cell body and overlap midcell (19, 20). These flagellar filaments are crucial for the flat-wave morphology of *B. burgdorferi* (21). Rotation of the periplasmic flagellum is driven by the motor, which acts as a rotary nanomachine at the base of each flagellum and plays a crucial role in flagellar assembly and directional switching (22–25). The flagellar motor is highly conserved among bacteria and composed of more than 25 different proteins (17), including those involved in the assembly of the MS ring, P-ring, and C-ring or switch complex, rod, and stator, respectively (15, 17, 26–29). Each stator complex is composed of two membrane proteins: MotA (5 copies) and MotB (2 copies). As stator complexes are recruited to the motor, they are powered by ion-motive force, thus generating torque for flagellar motility (11, 12, 30–32). FliL forms a ring around the stator complex, and interaction between the FliL ring and MotB is crucial to stabilize the stator complex in its extended state (33). In addition, the FliL-stator complex is further enclosed by flagellar collar proteins, suggesting that an ordered, cooperative assembly is necessary for the proper localization and stability of individual stator complexes (33–36). The collar and stator interact with each other and play crucial roles in stator assembly and/or stability, and the morphology and motility of *B. burgdorferi* (24, 36–38).

The collar is a large spirochete-specific multiprotein complex composed of an inner core and an outer turbine-like structure, although its composition and assembly are only partially understood. Using bioinformatics, genetics, biochemical analyses, and cryo-electron tomography (cryo-ET), we previously identified five collar proteins: FlbB, FlcA, FlcB, FlcC, and FlcD (35–39). FlbB appears to serve as the base of the collar structure, thought to provide bearing-like support for the MS-ring (36, 39). FlcD localizes near the base of the collar and contributes to overall collar assembly through interactions with FlbB (37). FlcA forms the turbine-like structure and directly interacts with the stator units (38), while FlcB constitutes the middle portion of the collar, and FlcC forms part of the upper region (35). The collar enables assembly of the flagellar motor within the highly curved membrane of spirochetes and serves as a structural scaffold for stator recruitment and stabilization. Importantly, we previously found that the collar proteins form extensive protein-protein networks with each other and with FliL and stator proteins MotA and MotB (39). However, the mechanisms by which the collar assembles and promotes stator assembly, processes that are critical for the unique motility of spirochetes, have remained unclear, partly because additional unidentified proteins are involved in collar assembly and function (39).

Using an integrated set of bioinformatics, structural, and functional analyses, we identify BB0298 as a collar protein adjacent to FlcA and FliL-stator complexes. Based on its peripheral localization within the collar, we designate BB0298 flagellar collar protein E (FlcE). Our studies suggest that FlcE functions as a latch that secures the FliL-stator complex, stabilizing the torque-generating units by tethering collar protein FlcA and FliL. This model is consistent with our finding that loss of *bb0298* reduces stator occupancy, resulting in cells with a nearly rod-shaped morphology and an almost non-motile phenotype. The presence of FlcE in motile spirochetes, along with conservation of its tetratricopeptide repeat (TPR) domains among other flagellar proteins across bacterial species (40, 41), suggests that the findings presented here have broad implications for understanding flagellar assembly and bacterial motility.

## Results

### Identification of BB0298 as a potential periplasmic flagellar protein

We previously identified five proteins, FlbB, FlcA, FlcB, FlcC, and FlcD, that contribute to the distinctive periplasmic flagellar collar structure of *B. burgdorferi* and are essential for flagellar orientation, stator assembly, cell morphology, and motility (35–39). However, substantial portions of the collar densities remained unassigned (39), indicating that additional proteins contribute to collar assembly. We therefore felt compelled to develop strategies to identify additional, previously unidentified periplasmic flagellar proteins in this spirochete. As illustrated in **Figure 1A**, our strategies identified BB0298 as a potential flagellar protein that met our criteria. Our recent studies suggested that some of the unidentified protein densities were around the periphery and could be membrane-binding proteins (39). Based on previous experimental analyses, BB0298 is indeed a periplasmic inner-membrane-binding protein (42–44). Furthermore, while BB0298 was annotated as a hypothetical protein of unknown function (45), it, like other periplasmic flagellar proteins, possesses a typical N-terminal leader peptide (43) that is cleaved by signal peptidase II after Ser15, while attaching a lipoprotein anchor to Cys16 (**Figure S1**). Moreover, BB0298 also contains tetratricopeptide repeat (TPR) domains (**Figure S2**), which are typically involved in extensive protein-protein interactions (46, 47) and have been identified in other flagellar proteins (35, 37, 38, 40, 41). Remarkably, the *bb0298* gene is located within the conserved bacterial *dcw* (*d*ivision and *c*ell *w*all) gene cluster, immediately preceded by *ftsW, ftsQ, ftsA, ftsZ* cell-division genes, followed by *dprA, hslVU,* plus a long flagellar operon that starts with *flgB* and includes 25 additional flagellar genes (**Figure S3**) (45, 48). Together, these observations strongly suggested that *bb0298* encodes a membrane-associated periplasmic flagellar protein.

**Figure 1.**
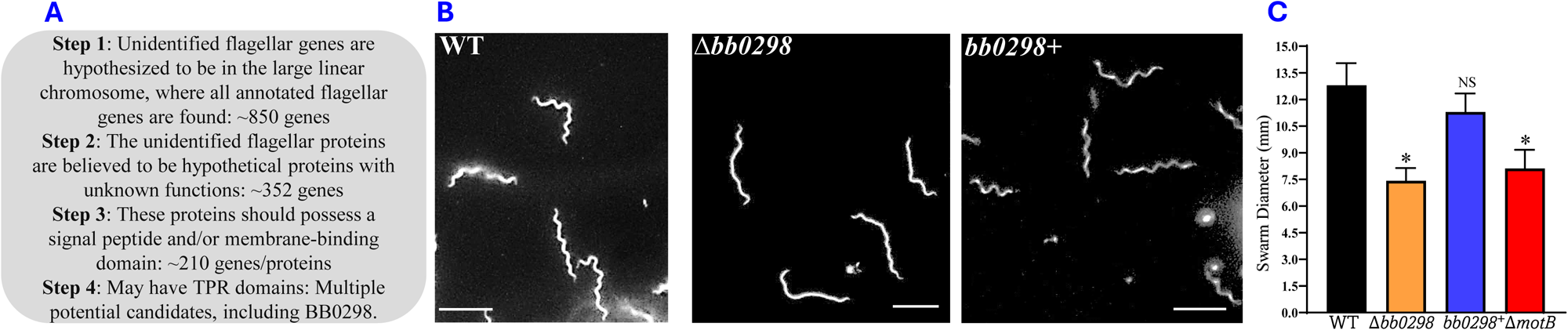
(**A**) Strategies used to determine potential new periplasmic flagellar proteins in *B. burgdorferi*. A total of 1,670 gene-encoded proteins were manually screened using the Spirochetes Genome Browser (http://sgb.leibniz-fli.de/cgi/list.pl?sid=24&c_sid=yes&ssi=free). This comprehensive analysis identified multiple potential candidates, including BB0298, hereafter referred to as FlcE. (**B**) Left three panels—Dark-field microscopy observations of the Δ*bb0298* or Δ*flcE* mutant cells display a nearly rod-shaped morphology, whereas the wild type or complemented *bb0298+* or *flcE+* cells exhibit the characteristic flat-wave morphology of the spirochete. The scale bar represents 10 μm. (**C**) Right panel—Swarm plate motility assays in 0.3% agarose mixed with BSK-II media indicate that Δ*flcE* mutant cells cannot swarm, similar to the completely non-motile Δ*motB* mutant cells. Complementation restored motility. An asterisk indicates a statistically significant difference between WT and mutant. NS, not significant

To specifically investigate the role of *bb0298* in flagellar motility, we inactivated this gene using a promoterless kanamycin resistance cassette (49) and confirmed the mutant clones by PCR (**Figure S4**). Subsequently, the Δ*bb0298* mutant cells were observed under a dark-field microscope. As shown in **Figure 1B**, the mutant cells exhibit altered morphology compared with the characteristic flat-wave morphology of the wild-type spirochete. Strikingly, the mutant cells exhibit a nearly complete loss of motility, as determined by swarm plate assays (**Figure 1C**). Although our gene-deletion strategy was specifically designed to minimize altering downstream gene expression (49), qRT-PCR analysis confirmed that the mutant exhibits no detectable polar effects (**Figure S5**). Even so, we complemented the Δ*bb0298* mutant with an intact copy of the gene transcribed from the flagellar *flgB* promoter (**Figure S4**) (18, 50, 51). Complemented *bb0298+* cells restored all phenotypes exhibited by the mutant, indicating that the *bb0298* deletion is responsible for the observed effects (**Figure 1 and other data below)**.

### BB0298 localizes to the collar periphery and affects periplasmic flagellar assembly

To understand the mechanism underlying the motility defect of the Δ*bb0298* mutant, we performed cryo-ET analyses of the periplasmic flagella in the Δ*bb0298* and *bb0298^+^* or wild-type cells. Although the mutant assembled flagella, some filaments were shortened and misoriented towards the cell pole rather than extending toward the opposite pole, as observed in complemented *bb0298^+^* [and wild-type cells (36–38)] (**Figure S6**). Similar defects in filament orientation have been reported in other collar mutants (36–38), suggesting that the altered morphology and motility defects of the Δ*bb0298* strain arise from abnormalities in the periplasmic flagella.

To further investigate the structure and function of BB0298, we employed subtomogram averaging to analyze flagellar motors from the Δ*bb0298* and *bb0298^+^* strains and compared their in-situ motor structures with those of wild-type (**Table S1**). Comparison of the Δ*bb0298* motor (**Figure 2A**) with the motors from the complemented (**Figure 2B**) and wild-type strains (**Figure 2C**) revealed the loss of a peripheral collar density, suggesting that BB0298 localizes to the outer edge of the collar (**Figure 2B-D**).

**Figure 2.**
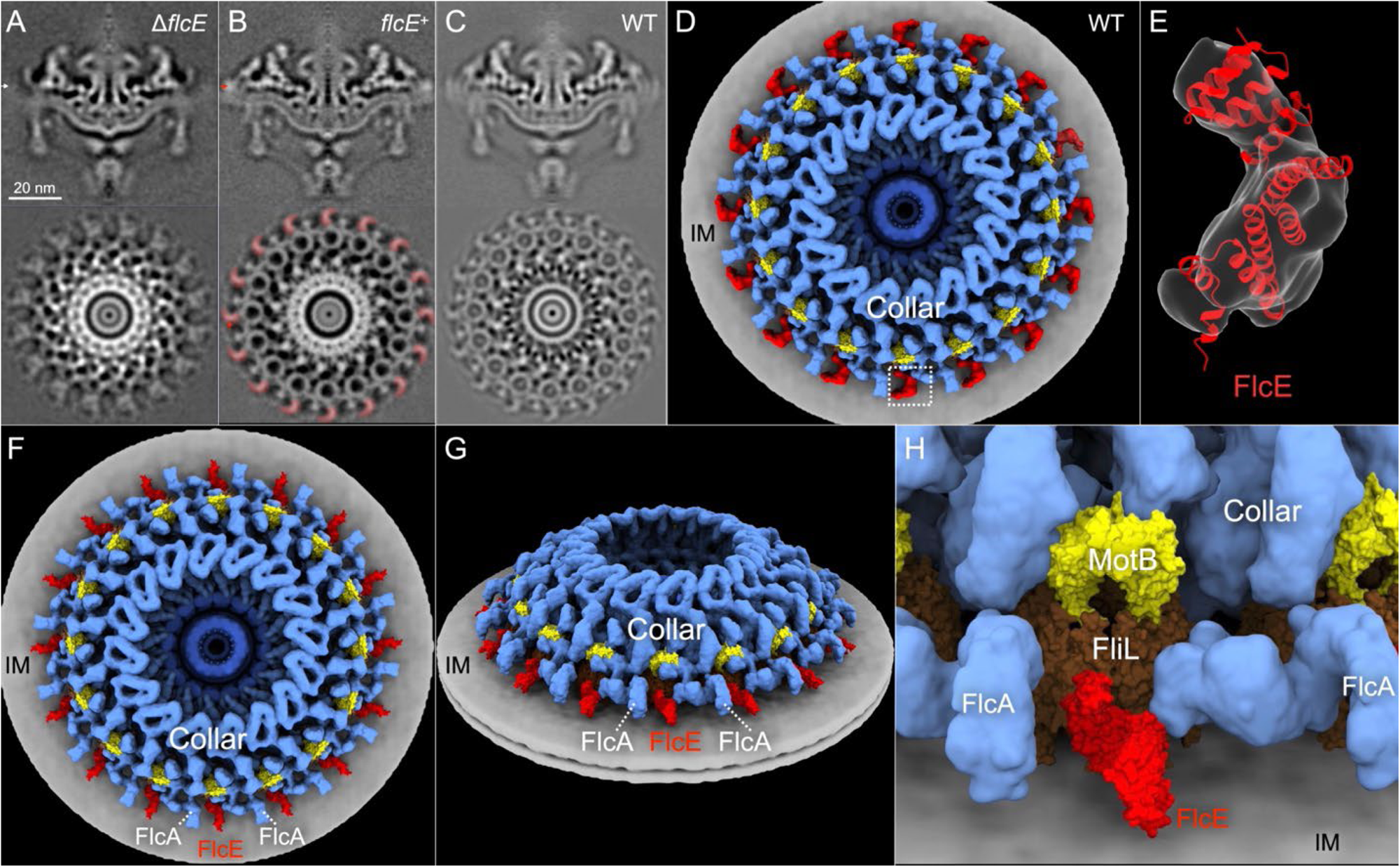
Comparative analysis of motor structures from Δ*flcE*, *flcE*+, and wild-type strains. (**A**) Two orthogonal sections from the subtomogram average of the motor in Δ*flcE*. (**B**) Two orthogonal sections from the subtomogram average of the motor in *flcE*+. Notably, both FliL-rings, the stator complexes, and the peripheral densities (highlighted in red) are absent in the Δ*flcE* but present in the *flcE+* motor. (**C**) Two orthogonal sections of the in situ structure of the wild-type motor. The FliL-rings, the stator, and peripheral densities are also present in these motors, as in complemented cells. (**D**) Top view of the surface rendering of the wild-type motor structure with the additional peripheral densities colored red and stator units in yellow. (**E**) Zoom-in of the AlphaFold-predicted FlcE model, fitted into each peripheral density. (**F, G**) Top and tilted views of the wild-type motor structure showing that the FlcE model occupies the gap between adjacent FlcA monomers (light blue). (**H**) Zoomed-in view illustrating that FlcE effectively encloses the FliL (brown)-stator complex (yellow).

To test this interpretation, we used AlphaFold3 to predict the structure of BB0298 and fitted the model into the density present in the wild-type and complemented *bb0298+* motors, but absent in the deletion mutant. The predicted structure matched the peripheral collar density well, with a cross-correlation of ∼0.7 calculated in ChimeraX (**Figure 2E**), indicating that BB0298 contributes at least in part to this peripheral density. Based on its localization and function, we renamed BB0298 as FlcE, consistent with the nomenclature of previously identified collar proteins.

The modeled FlcE structure localizes adjacent to the 16-solenoid, S-shaped FlcA structure that we recently described (39). Notably, FlcE occupies a position between neighboring FlcA subunits (**Figure 2F-H**), suggesting direct interaction between the two proteins. To experimentally test this model, we performed pull-down assays and far-Western blotting. Both assays demonstrated that FlcE specifically interacts with FlcA among the collar and flagellar proteins tested (**Figure 3A-D**).

**Figure 3.**
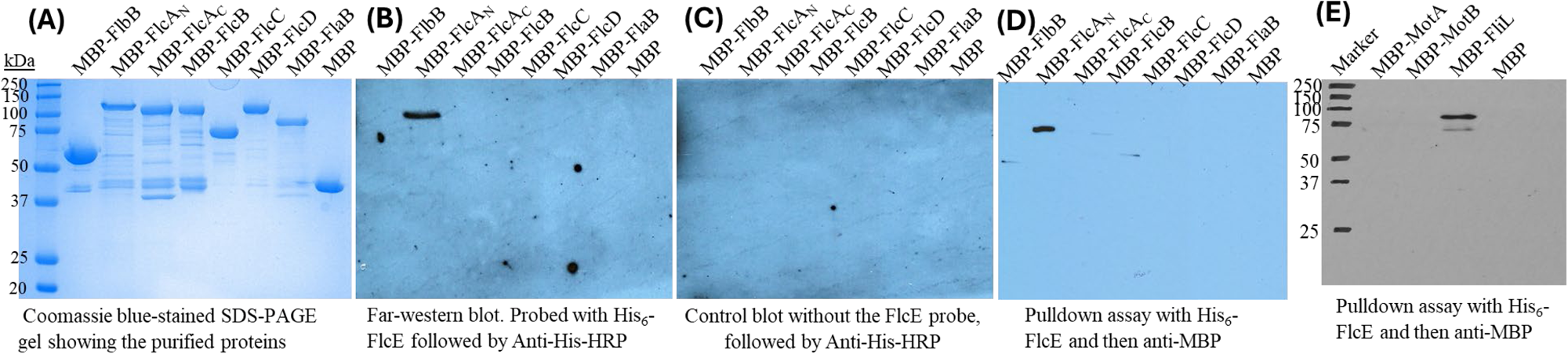
Analysis of FlcE interactions with FlcA and FliL by far-western blotting (A-C) and pull-down assays (D-E). One microgram of each purified recombinant protein was subjected to SDS-PAGE and analyzed by Coomassie blue staining (**A**) or Far-Western blotting using the His6-FlcE/BB0298 probe (**B**), alongside a control blot (**C**). Pull-down assays showed that FlcE interacts only with the N-terminus of FlcAaa2-328 (**D**) and FliL, not with MotA or MotB (**E**).

Previous work showed that FlcA binds the stator protein MotB through its C-terminal domain (residues 360–931) (38). To identify the region responsible for interaction with FlcE, we performed those assays using FlcA truncations. Both these experiments showed that FlcE interacts specifically with the N-terminal region of FlcA (residues 2-328) (**Figure 3B-D**). Together, these findings indicate that FlcA uses distinct domains to bind MotB and FlcE, thereby linking the collar periphery to the stator complex.

### FlcE restrains the FliL-stator ring complexes

*B. burgdorferi* flagellar motors normally assemble 16 FliL-stator complexes surrounding the rotor (33, 35). A focused classification of the stator complex densities was performed to quantify FliL-stator occupancy, as previously described (24). In the Δ*flcE* mutant, one class average retained assembled stator complexes, as indicated by MotA density, whereas another class lacked them entirely (**Figure 4A, B**). By contrast, complemented *flcE*+ cells restored intact stator assembly (**Figure 4C**). Loss of FlcE reduced stator occupancy from ∼98% in wild-type motors to ∼75%, corresponding to the loss of approximately four stator units per motor (**Figure 5**). Immunoblot analyses showed that FliL and MotB protein levels were unchanged (**Figure S7**), indicating that the defect results from impaired stator stabilization rather than reduced protein synthesis. Despite retaining relatively high (∼75%) stator occupancy, the Δ*flcE* mutant was nearly non-motile (**Figure 1C**), demonstrating that near-complete assembly and stabilization of the FliL-stator ring are essential for robust spirochetal motility (**Figure 5**).

**Figure 4.**
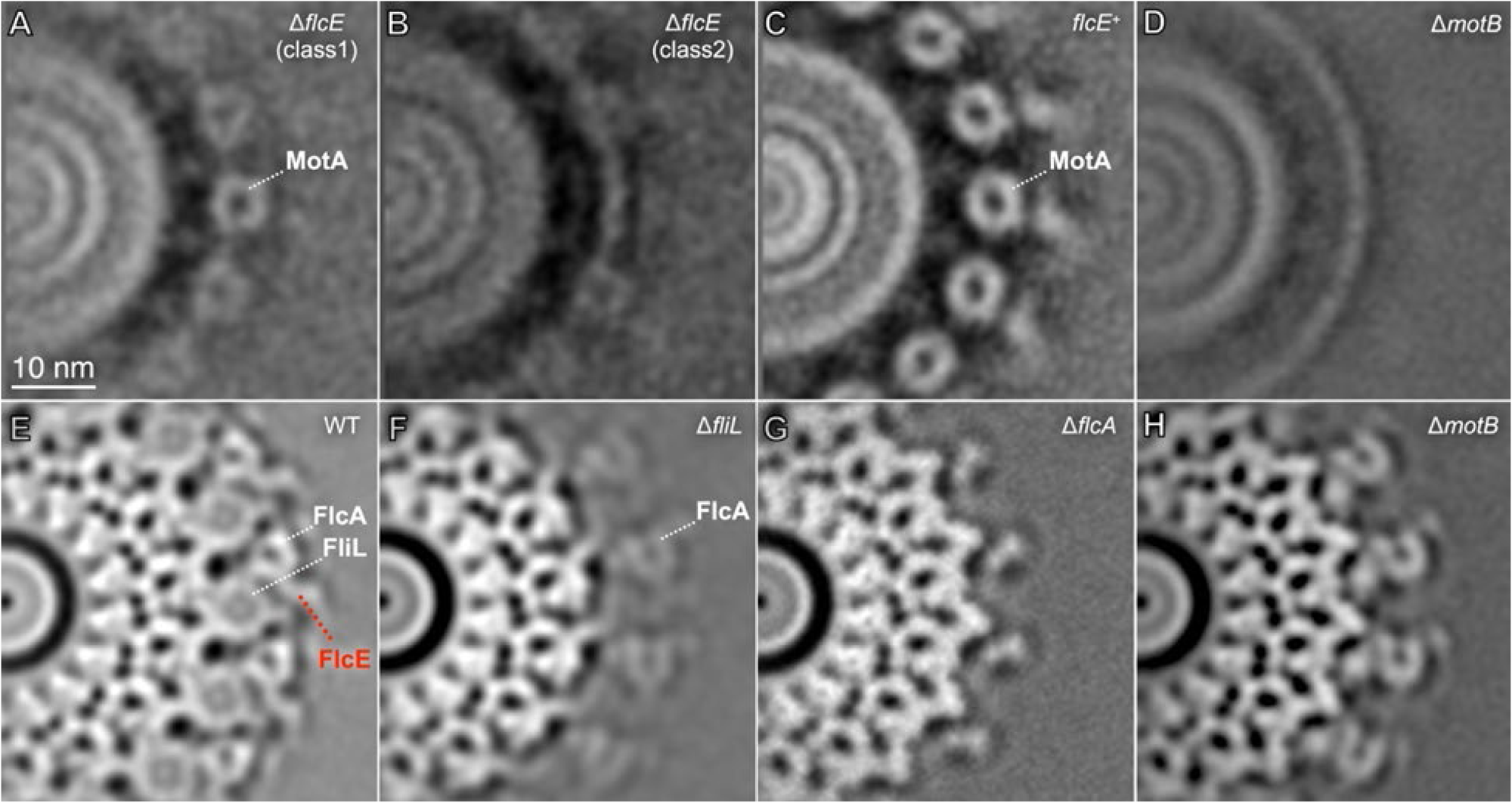
Comparison of motor structures from Δ*flcE*, *flcE*+, wild type (WT), Δ*fliL*, Δ*motB,* and Δ*flcA*. (**A**) Section from a class average of the Δ*flcE* motor showing a visible MotA complex. (**B**) Section from another Δ*flcE* class average in which the MotA complex is absent. (**C**) Section from the subtomogram average of the complemented *flcE*+ motor showing a clearly resolved MotA complex. (**D**) Section from the Δ*motB* motor in which the MotA complex is absent. (**E**) Section from the WT motor subtomogram average, displaying the FliL-ring, FlcA, and FlcE. (**F**) Section from the Δ*fliL* motor subtomogram average showing FlcA but lacking FlcE. The FlcA density appears relatively weak, suggesting increased flexibility in the absence of FliL. (**G**) Section from the Δ*flcA* motor subtomogram average, which lacks FlcA, FliL, and FlcE. (**H**) Section from the Δ*motB* motor in which the FliL-ring and FlcE are absent.

**Figure 5.**
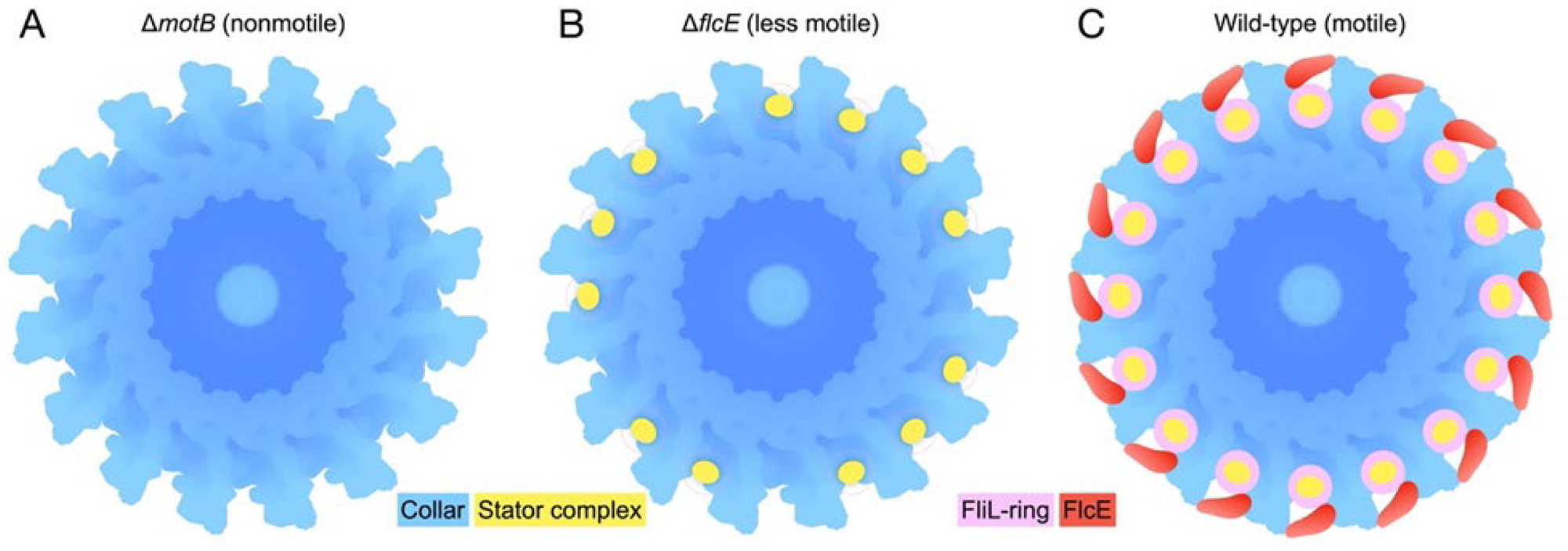
Flagellar motor models in Δ*motB,* Δ*motB,* and wild-type. (**A**) Δ*motB* mutant cells are deficient in these stator units and exhibit a nonmotile phenotype. (**B**) Δ*flcE* mutants can assemble approximately 75% of stator units but have significantly reduced motility. (**C**) Wild-type spirochetes possess functional motors and FlcE, which latch the FliL-stator units, enabling efficient swimming through the complex environments of ticks and mice, highlighting their remarkable adaptability and persistence.

We also reanalyzed previously reported motor mutants to determine whether the density assigned to FlcE depends on other collar-stator components. The FlcE density was absent in Δ*motB*, Δ*fliL*, and Δ*flcA* mutants (**Figure 4D-H**)(24, 33, 38). Consistent with earlier studies, stator density was also lost in the Δ*flcA* and Δ*fliL* motors, indicating that disruption of the FliL-stator assembly destabilizes FlcE localization. Together, these findings demonstrate that FlcE, FlcA, and the FliL-stator complexes are structurally interdependent.

Cryo-ET structures further revealed that FlcE aligns directly with the 16 FliL-stator complexes (**Figure 2**). The crescent-shaped FlcE appears to wrap around the FliL-stator assembly like a “seatbelt,” tethering FlcA on one side and FliL on the other (**Figure 5**). Consistent with this model, protein-protein interaction assays confirmed direct binding between FlcE and FlcA, as well as between FlcE and FliL, whereas no interaction was detected with MotA or MotB (**Figure 3E**). Together, these findings support a model in which FlcE functions as a molecular latch that restrains and stabilizes the torque-generating FliL-stator complexes, thereby preserving motor integrity and enabling efficient motility.

### The TPR repeats of FlcE are conserved in other bacteria

Sequence analysis of FlcE (BB0298) revealed that it is well conserved among members of the spirochetal orders Spirochaetales and Winmispirales (**Figure S1)**, including the syphilis pathogen *Treponema pallidum* and oral pathogen *Treponema denticola*, which is widely used as a model organism to study spirochete physiology. BB0298 orthologs are seen in motile spirochetes from the families *Borreliaceae, Breznakiellaceae*, *Spirochaetaceae, Treponemataceae*, and *Winmispiraceae*, but are absent from motile members of the *Leptospiraceae* and non-motile members of the *Sphaerochaetaceae* (**Table S2**). These proteins contain similarly positioned predicted TPR domains (**Figure S1**) and are encoded within conserved genomic neighborhoods (**Figure S3 and Table S2**). TPR domains are well known for mediating protein-protein interactions (46, 47). Importantly, several bacterial TPR-containing proteins have been proposed to participate in flagellar biogenesis (40, 41). Although the amino acid sequences of TPR-containing proteins such as FlcE are highly divergent, complicating identification of homologs, our findings, together with previous studies, support the idea that many bacterial TPR-containing proteins are involved in flagella-associated functions (37, 40, 41, 52–55).

### FlcE fills a missing link within the collar-stator-rotor interaction network

Our recent study revealed the periplasmic flagellar collar structure and its interactions with the FliL-stator rings and the MS ring (39). However, that previous model contains substantial unassigned collar densities around the FliL-stator rings. Here, we provided evidence that FlcE contributes to the peripheral region of the collar (**Figures 2, 4, 5**). Importantly, FlcE, together with the other identified collar-stator proteins, forms an extensive collar-stator-rotor interaction network, as supported by cryo-ET and genetic analyses, structural modeling, and experimentally validated protein-protein interactions (39). Interactions among FlcE, FliL, and FlcA introduce additional structural connectivity (**Figure S8B**), substantially improving our understanding of how the periplasmic flagellar motor, particularly the collar-stator, is organized to generate the torque required for Lyme disease spirochetes to navigate complex tick-vertebrate host environments (56). To fully understand the intact collar-stator architecture and its function at the molecular level, additional proteins remain to be identified to account for unassigned densities in the collar (**Figure S8A**).

## Discussion

Identifying the proteins responsible for the spirochete-specific collar in the *B. burgdorferi* flagellar motor has been a longstanding challenge because none of the initially annotated flagellar genes encoded collar proteins (45). FlcE identified here occupies part of the peripheral collar density and expands the known collar architecture to six proteins (35–39). These findings strengthen the emerging view that the spirochetal collar is a specialized, multiprotein scaffold that stabilizes the torque-generating stator complexes required for motility and virulence (2, 3, 35–39).

Our data support a model in which FlcE functions as a latch stabilizing the FliL-stator complexes (**Figures 2 and 5**). Loss of FlcE abolishes the FliL ring, reduces stator occupancy, and severely impairs motility despite normal expression of FliL and MotB (**Figures 1, 4, 5, and S7**), indicating that FlcE primarily stabilizes assembly, rather than synthesis, of the engaged stator complex. Structural modeling and interaction studies further show that FlcE connects collar protein FlcA to the FliL-stator complex (**Figure 3**) (24, 33), providing a direct mechanistic explanation for how the collar reinforces high-torque stator engagement. Without this peripheral restraint, stator complexes would likely disengage, tilt, or fail to maintain the geometry required for efficient torque transmission (**Figure 5**).

Our findings also refine the functional organization of the collar. FlcA appears to act as a central interaction hub, using distinct domains to bind MotB (38) and FlcE simultaneously, thereby integrating the collar with the stator complex (**Figures 2, 3**). This extensive connectivity suggests that collar integrity and stator engagement are cooperatively linked, explaining why even partial reductions in stator occupancy produce dramatic motility defects in spirochetes (**Figures 1 and 5**).

Importantly, the conservation of TPR-containing FlcE-like proteins in diverse flagellated bacteria suggests that peripheral scaffolding of stator complexes represents a broad evolutionary strategy for sustaining high torque generation. Because TPR-containing proteins are often structurally conserved despite low sequence similarity, FlcE also highlights the limitations of homology-based annotation in identifying specialized motility factors (40, 41).

Together, our results establish FlcE as a critical structural component of the collar-stator-rotor interface and provide a mechanistic framework for understanding how spirochetes stabilize the exceptionally high-torque motors (56, 57) required for motility within complex tick-vertebrate host environments. At the same time, additional collar proteins remain to be discovered for the intact molecular architecture of the spirochetal motor to be fully resolved.

## Materials and methods

### Bacterial strains and growth conditions

The wild-type (WT) clone used in this study was the high-passage *B. burgdorferi* strain B31-A. (51, 58). The Δ*bb0298* mutant and its complement strains were constructed as described below. *B. burgdorferi* cells were grown in liquid Barbour-Stoenner-Kelly (BSK-II) medium or agarose plates and incubated at 35°C in a 2.5% CO2 incubator, as described previously (49, 59). Antibiotics, when needed, were added to the *B. burgdorferi* medium at the following concentrations: 200 μg/ml of kanamycin and/or 100 μg/ml streptomycin. *Escherichia coli* strains were cultured at room temperature or 37°C in liquid Luria-Bertani broth or plated on LB agar (60). When needed, 100 μg/ml ampicillin, 100 μg/ml spectinomycin, 0.2% glucose, 80 μg/ml 5-bromo-4-chloro-3-indolyl-β-d-galactopyranoside (X-Gal), and/or 0.5 mM isopropyl-β-d-1-thiogalactopyranoside (IPTG) were added to LB medium as supplements.

### Construction of *bb0298* mutants and complemented strains

Construction of the *bb0298* inactivation plasmids, electroporation, and plating conditions were described previously (49, 59). Briefly, a 300 bp upstream sequence along with the *bb0298* orf DNA (993 bp) was PCR-amplified using Q5® High Fidelity DNA Polymerase with primers (5’-3’) FP-BB0298-Upper PI-Kan (ACTGTAATATATGGTCATGCTATTAATTCGAATCTTG) and RP1-BB0298-PI-KO (TTACTTCAAATTACTCATGATCCTAGAAACACCTTCT and then the PCR product was ligated into the pGEM-T Easy vector (Promega), yielding pGEM-T Easy::bb0298. The Pl-Kan (49) was similarly PCR-amplified from P_flgB_-Kan (51) using primers (5’-3’) PrLs Kan Hind-F (AAGCTTTAGTTAAAAGCAAT) and PrLs Kan Hind-R (AAGCTTTTAGAAAAACTCAT). The HindIII-restricted Pl-Kan DNA was then inserted into the HindIII sites within bb0298 (located 240 and 1092 base pairs from the bb0298 ATG start codon) of the pGEM-T Easy::bb0298 vector, yielding pGEM-T Easy::bb0298-Pl-Kan. This insertion mutation was in-frame. The direction of transcription of Pl-Kan was checked by another PCR using primers (5’-3’) FP-BB0298-Upper PI-Kan (ACTGTAATATATGGTCATGCTATTAATTCGAATCTTG) and RP2-BB0298-PI-KO-HindIII (GTACTGACCAAGCTTTTAGAAAAACTCATCGAGCATC). bb0298-Pl-Kan was digested and separated from pGEM-T Easy::bb0298-Pl-Kan vector by NotI restriction enzyme, and then the linear DNA was prepared for electroporation into strain B31-A competent cells. The transformants were selected with 200 μg/ml kanamycin. The kanamycin-resistant transformants were isolated, and the insertion of Pl-Kan was confirmed by PCR.

The Δ*bb0298* mutant was complemented *in trans* using the widely used shuttle vector pBSV2G, in which the *flgB* promoter is fused to an intact copy of the *bb0298* gene, as described (18, 61). Briefly, the pGEMT::PflgB-bb0259 (62) is used as a template to amplify BamHI-PflgB, and the bb0298-PstI DNA fragment was amplified from the B31A genomic DNA. The *flgB* promoter is then fused to the *bb0298* by NEBuilder® HiFi DNA Assembly kit and subsequently ligated into the pGEM-T Easy vector (Promega Inc.), yielding plasmid pGEMT::PflgB-bb0298-Easy. The plasmid containing the PflgB-bb0298 fragment was then released using BamHI and PstI. The shuttle vector pBSV2G was similarly prepared and ligated, generating pBSV2G::PflgB-bb0298. The vector pBSV2G::PflgB-bb0298 DNA was then electroporated into the Δ*bb0298* mutant cells, followed by selection with both kanamycin and gentamicin. The resistant clones were analyzed by PCR using appropriate primers.

### Bioinformatics

The PSI-BLAST search tool and HHPred (63, 64) were used to identify BB0298 homologs in protein sequence databases. Transmembrane domains and signal peptides were predicted using DeepTMHMM and SignalP 6.0 programs (65, 66). TPR domain predictions and genomic neighborhoods were taken from the RefSeq database (67). Protein domains were analyzed using the Conserved Domain Database and Pfam (68, 69).

### Dark-field microscopy and swarm plate assays

Exponentially growing *B. burgdorferi* cells were observed using a Zeiss Axio Imager M1 dark-field microscope equipped with an AxioCam digital camera to assess morphology and motility, as described previously (59, 70). A swarm plate assay was performed to determine the motility of spirochetes using our established protocol (57, 68) with exponentially growing *B. burgdorferi* cells. Data is shown as the mean of three assays with standard deviation, and a Student’s t-test was performed to assess significance, with P-values under 0.05 indicating significance.

### Overexpression of recombinant proteins in *E. coli*

To express the *B. burgdorferi* BB0298 protein in *E. coli*, a DNA fragment harboring the BB0298 open reading frame (ORF) without the signal peptide region (1 to 20 amino acids) was PCR amplified from chromosomal DNA of *B. burgdorferi* B31-A cells using primers R-0298-BamHI F (GGTCGC<u>GGATCC</u>AAAGAAAAATCAAATCTTGGTCT) and R-0298-NotI R (TCGAGT<u>GCGGCCGC</u>CTTCAAATTACTCATGATCCT) (restriction enzyme sites are underlined) and cloned into the pET28a(+) (Novagen Inc.) using BamHI and NotI restriction sites. Expression of MBP, MBP-FliL, MBP-FlbB, MBP-FlaB, MBP-FlcB, MBP-FlcC, MBP-FlcD, MBP-FlcA_C (aa 360-931)_, MBP-FlcA_N (aa 2-328)_, MBP-FliL, MBP-MotA, and MBP-MotB was described elsewhere (24, 33, 35–39). All *E. coli* strains were induced with 0.5 mM IPTG at room temperature, and the purification of recombinant proteins was carried out using amylose resin for MBP-tagged proteins and HisPure Ni-Nitrilotriacetic Acid (NTA) resin for His-tagged proteins.

### SDS-PAGE, immunoblot, and far-western blotting

Sodium dodecyl sulfate polyacrylamide gel electrophoresis (SDS-PAGE) and immunoblotting with enhanced chemiluminescent detection were performed as described (GE Health Inc) (3). Protein concentrations were determined using a Bio-Rad protein assay kit with bovine serum albumin as the standard. Unless specified otherwise, 10 μg of protein was subjected to SDS-PAGE and immunoblotted with specific antisera, as described (3).

The interactions of two recombinant proteins were determined by Far-Western blotting or affinity blotting, as described previously (71). Briefly, 1 µg of each purified recombinant protein was subjected to SDS-PAGE and subsequently transferred to polyvinylidene difluoride (PVDF) membranes. A parallel SDS-PAGE gel was stained with Coomassie Brilliant Blue. The PVDF membranes were blocked with 5% skim milk, 10 mM Tris, 150 mM NaCl, and 0.3% Tween 20 (pH 7.4) for 4 to 6 hours at room temperature. The membranes were then incubated overnight at 4°C with or without purified recombinant His_6_-BB0298 protein at 2 µg/ml in the blocking solution. After three washes with a buffer containing 10 mM Tris, 150 mM NaCl, and 0.3% Tween 20 (pH 7.4), the membranes were probed with the monoclonal anti-His-HRP® M2 antibody (Sigma-Aldrich Co. LLC).

### Pull-down assays

Pull-down assays were used to determine interactions between two proteins, as described (39). Briefly, expression vectors with cloned *B*. *burgdorferi* gene coding for MotA, MotB, FliL, or FlcE/BB0298 were described above and previously. The *E*. *coli* cells harboring the expression plasmids were induced separately per the manufacturer’s instructions. The expressed cells were mixed (∼1:1; e.g., BB0298-His_6_ and MBP-MotA, MBP-MotB, or MBP-FliL), French-pressed to lyse the bacteria, and purified using Ni-NTA agarose resin. The purified proteins were subjected to SDS-PAGE and then transferred to a PVDF membrane for immunoblotting with a commercially available Anti-MBP-HRP monoclonal antibody (Invitrogen Inc.) according to the manufacturer’s instructions. The pull-down experiments were conducted twice to ensure scientific rigor and reproducibility.

### Sample preparation for cryo-ET

*B. burgdorferi* cells in liquid were resuspended with phosphate buffer saline (PBS) to a final OD_600_ of 1.0. 10 nm of gold tracer solution (Aurion) was then added to the bacterial sample, and 5 μL of the mixture was deposited on discharged cryo-EM grids (Quantifoil). The front of the grids was blotted with filter paper (WhatmanTM), and the grids were then plunged into a liquid ethane and propane mixture using a Leica EM GP2 plunger. GP2 environmental chambers were set to 25 °C and 95% humidity. The grids were blotted from the front for 6s.

### Cryo-ET imaging

Frozen-hydrated specimens of *B. burgdorferi* were imaged below −180°C using Titan Krios G2 300kV transmission electron microscope (ThermoFisher) equipped with a field emission gun, K3 direct detection camera (Gatan), and GIF BioQuantum K3 Imaging Filter (Gatan). SerialEM software (72) was used to record images at 42,000x magnification with a physical pixel size of 2.148Å. The defocus was set as 4.8μm. To collect tilt series, the stage in the microscope was tilted from −48° to +48° in increments of 3° using the dose-symmetric scheme in FastTomo script (73, 74). The total electron dose was almost 70e^-^/Å^2^ distributed across 33 images in the tilt series. The dose fraction mode in SerialEM (72) was used to record the images.

### Cryo-ET data processing

MotionCor2 (75) was used to correct image drifting induced by the electron beam during image recording. IMOD software was used to create image stacks and then track with fiducial beads to align all images in each tilt series (76, 77). Gctf (78) was used to estimate defocus for all aligned images, and the contrast transfer function (CTF) was corrected using the ctfphaseflip function in IMOD (79). To reconstruct 4x binned tomograms, 4x binning of images in the tilt series was created using the binvol function in IMOD. Tomo3D (80) was then used to reconstruct tomograms. In total, we got 59 tomograms from Δ*bb0298* mutant cells and 50 tomograms from *bb0298^+^* complemented cells (**Table S1**). Dragonfly (Comet Technologies Canada Inc.) was utilized to segment the bacterial features, including the outer membrane, inner membrane, and flagellar filaments of varying lengths. In-situ structures of the flagellar motors were determined by using I3 (81, 82), as described previously (17, 33, 39).

## Acknowledgments

We thank Jorge Benach and James Carroll for sharing DnaK and MotB antibodies. We thank Jennifer Aronson for critical reading of the manuscript. M.Y.G. was supported by the Intramural Research Program of the National Institutes of Health. The contributions of the NIH authors are considered Works of the United States Government. The findings and conclusions presented in this paper are those of the authors and do not necessarily reflect the views of the NIH or the U.S. Department of Health and Human Services.

## Supplement Figures

**Figure S1.**
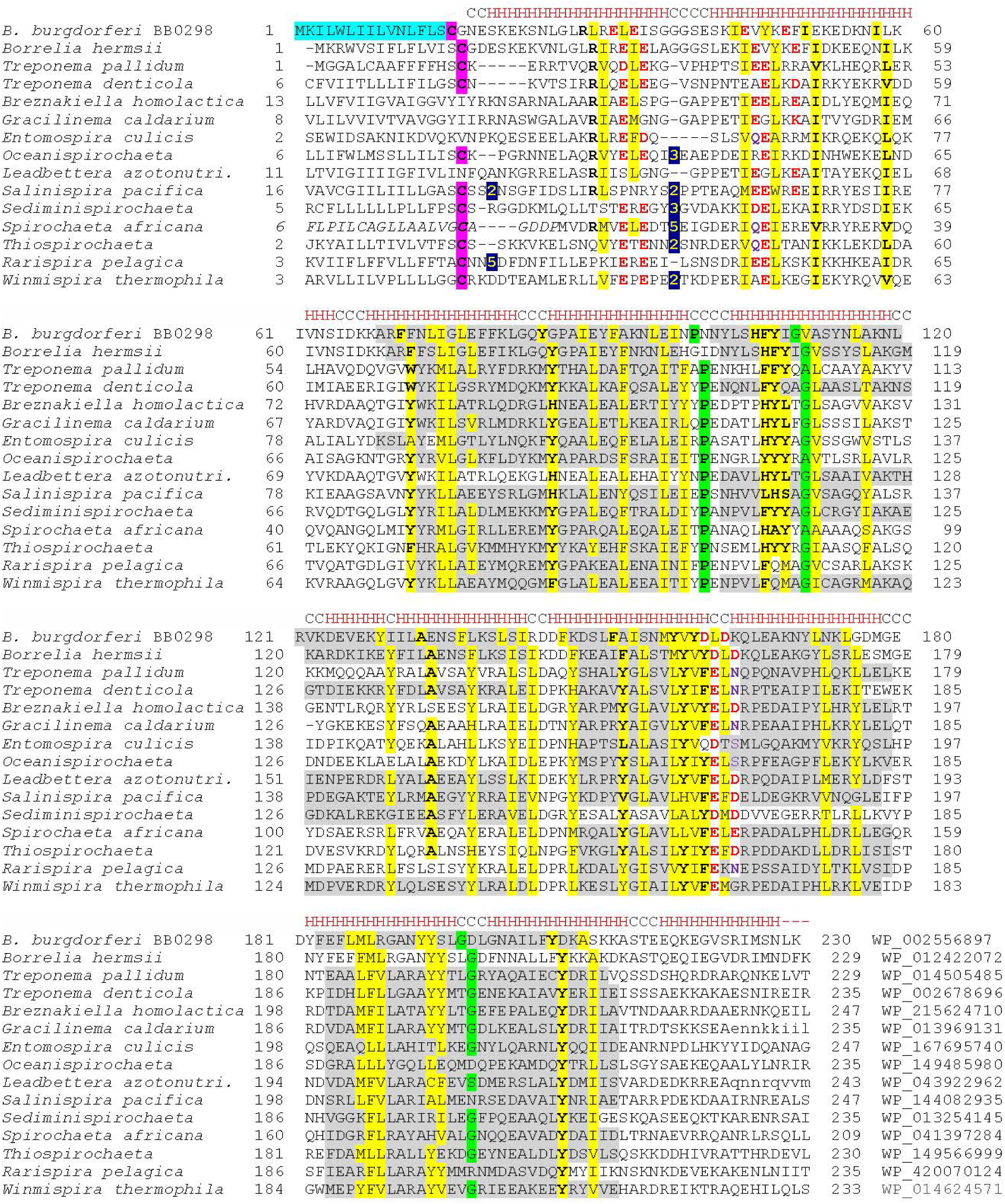
Sequence alignment of BB0298 and its homologs. The first line in each block represents the predicted secondary structure of BB0298 (C, coil; H, α-helix). Predicted leader peptide of BB0298 (PSort motif PS51257) is shaded blue, and the lipid attachment Cys residue is highlighted in magenta. The numbers represent amino acid positions and the lengths of the gaps between the aligned blocks. Conserved residues are in bold and colored red (Asp, Glu) or purple (Gln, Asn). Conserved hydrophobic residues are shaded yellow, turn residues (Pro, Gly) are shaded green. The sequences’ source organisms and locus tags (top to bottom) are as follows: *Borrelia burgdorferi* B31, BB0298; *Borrelia hermsii* DAH, BH0298; *Treponema pallidum* str. Nichols, TPANIC_0392 (TP_0392); *Treponema denticola* ATCC 35405, TDE_1206; *Breznakiella homolactica* RmG30, JFL75_10565; *Gracilinema caldarium* DSM 7334, Spica_1679; *Entomospira culicis* BR151, PVA46_05430; *Oceanispirochaeta crateris* K2, EXM22_07825; *Leadbettera azotonutricia* ZAS-9, TREAZ_RS06765; *Salinispira pacifica* DSM 27196, L21SP2_1167; *Sediminispirochaeta smaragdinae* DSM 11293, Spirs_1554; *Spirochaeta africana* DSM 8902, Spiaf_2113 (with corrected start, added residues in italics); *Thiospirochaeta perfilievii* DSM 19205, EW093_03175, *Rarispira pelagic*a 38H-sp 4, WKV44_08975; and *Winmispira thermophila* DSM 6578, Spith_0926. The last column shows the corresponding RefSeq accession numbers. TPR domain predictions, taken directly from the respective RefSeq entries, are indicated with grey shading

**Figure S2.**
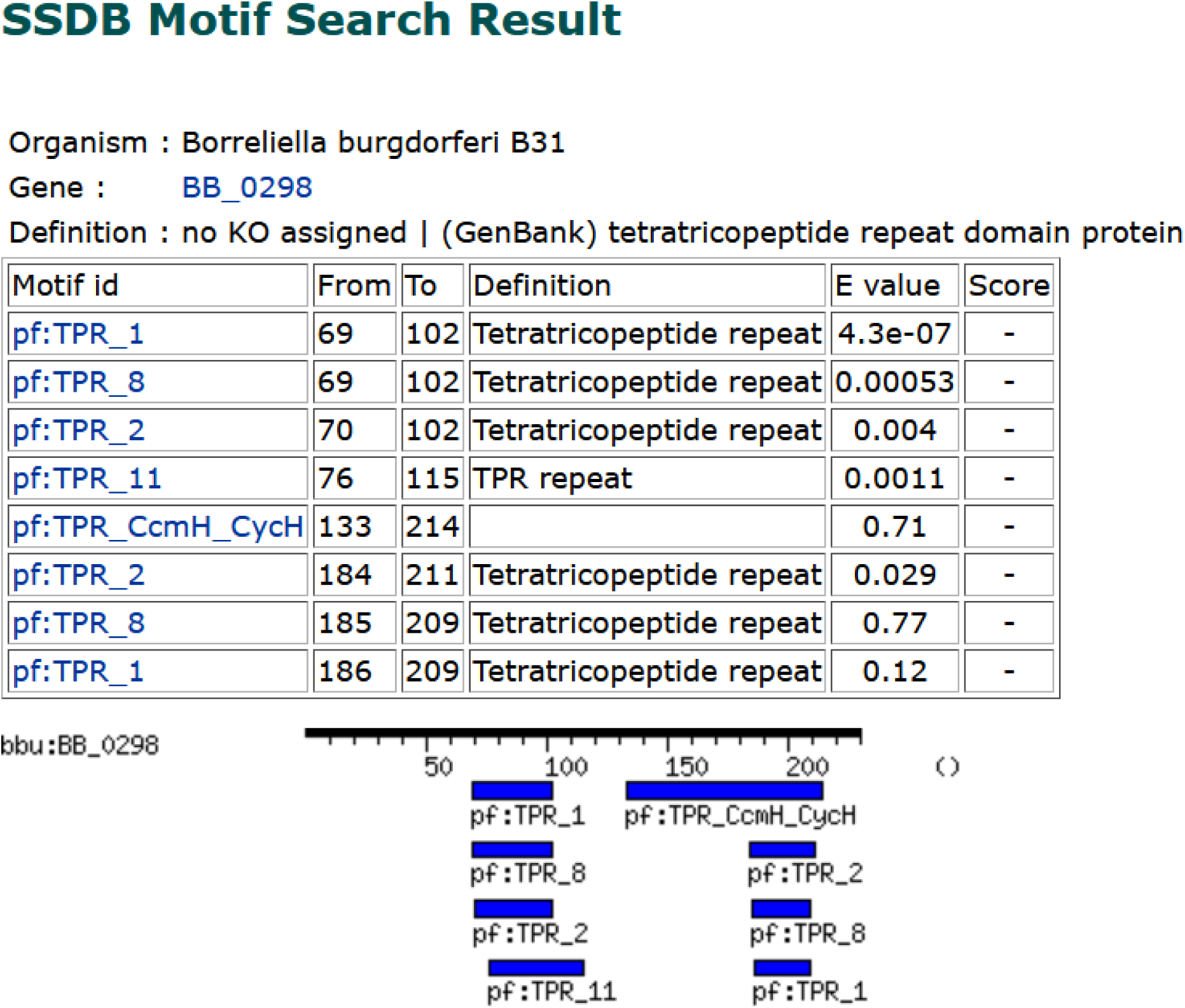
BB0298 or BB_0298 features tetratricopeptide repeat (TPR) domains. The motif search results generated from the SSDB using the KEGG genome browser for the *Borrelia burgdorferi* B31 genome are presented here. Each motif is provided with its length, the amino acid sequence position in BB0298, and the corresponding E-value. The lower the E-value, or the closer it is to zero, the more significant the match is.

**Figure S7.**
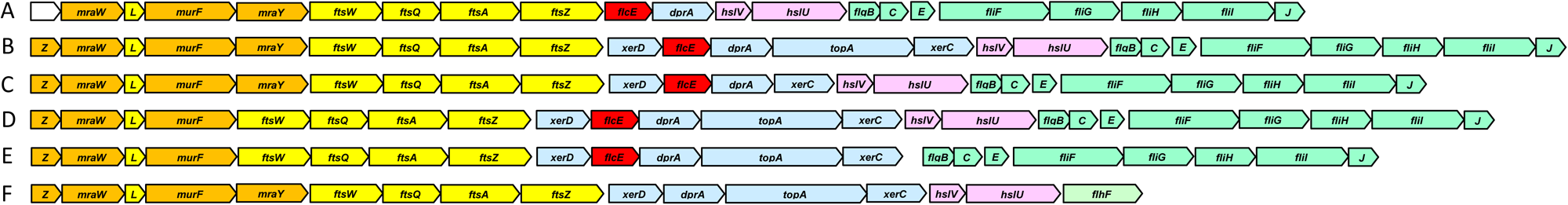
Organization of the *dcw* (*d*ivision and *c*ell *w*all) gene cluster in *Borrelia* and other spirochete genomes. Cell wall genes are orange, cell division genes are in yellow, DNA processing genes are in blue, ATP-dependent protease subunits are in pink, flagellar genes are in green, *flcE* (BB_0289) orthologs are in red. Abbreviated gene names are as follows: *Z*, *mraZ*; *C*, *flgC*; *E*, *fliE*; *J*, *fliJ*. The operons are generally conserved at the family level and are representative of the following lineages: A, *Borreliaceae* (represented by *Borrelia burgdorferi*, GenBank entry AE000783.1: positions 316,143..297,038); B, *Breznakiaceae* and *Winmispiraceae* (*Breznakiella homolactica,:* CP067089.2: 477,555..499,882); C*, Spirochaetaceae* (*Sediminispirochaeta smaragdinae,* CP002116.1: 1,661,125..1,682,188); D,E, *Treponemataceae* (D, *Treponema denticola,* AE017226.1: 1,230,092..1,252,475; E. *Treponema pallidum,* AE000520.1: 408,169..428,859); F, *Sphaerochaetaceae* (*Parasphaerochaeta coccoides,* CP002659.1: 759,739..744,520). See Supplementary Table S2 for details.

**Figure S4.**
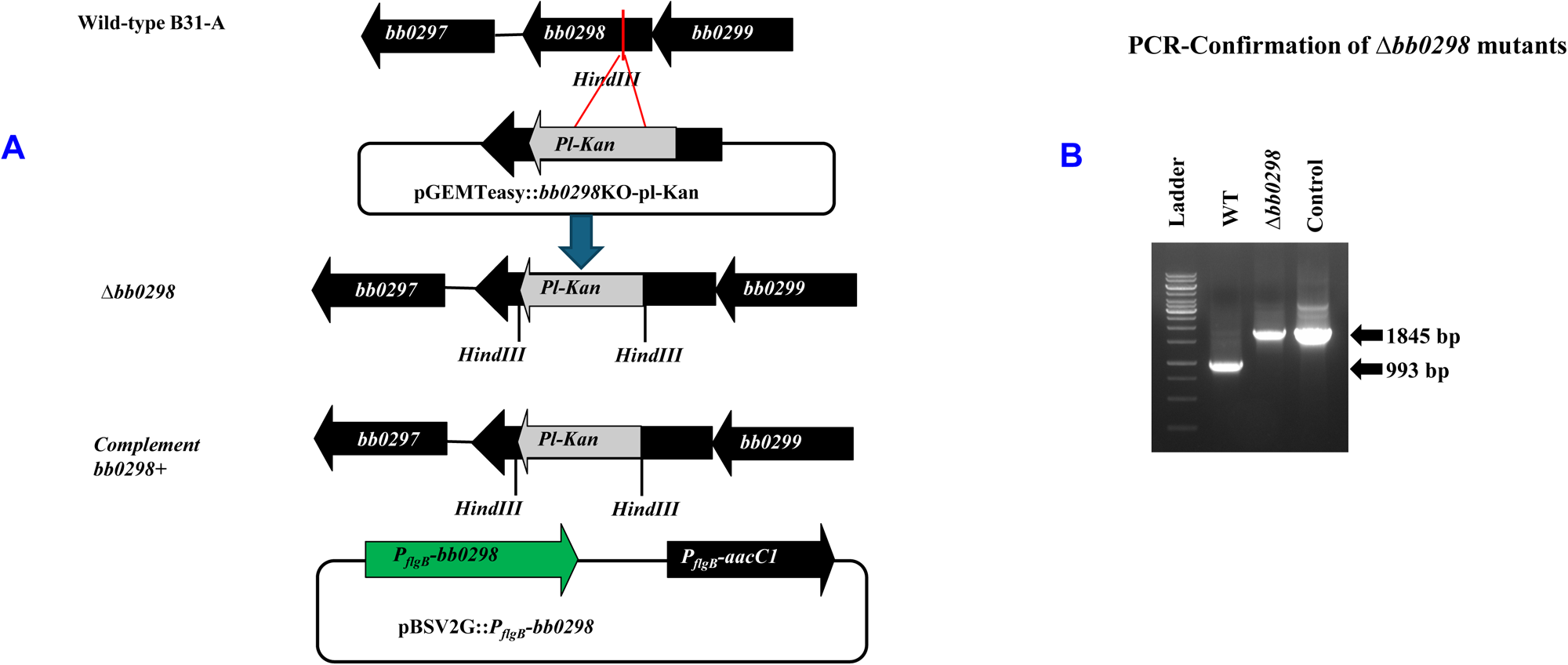
Inactivation of *bb0298* and complementation of the Δ*bb0298* mutant in *B. burgdorferi.* (A) In-frame insertion of the Pl-Kan cassette within the *bb00298* gene by homologous recombination resulted in the creation of the Δ*bb0298* mutant. The approximate location of Pl-Kan is shown by a thick grey arrow within the *bb0298* gene. (B) The integration of Pl-Kan within the *bb0298* gene was confirmed by using PCR; the control represents the input pGEMTeasy::bb0298KO-pl-Kan plasmid used to construct the mutant. Shuttle vector pBSV2G::PflgB-bb0298 was used to complement the mutant as described in the Methods

**Figure S5A.**
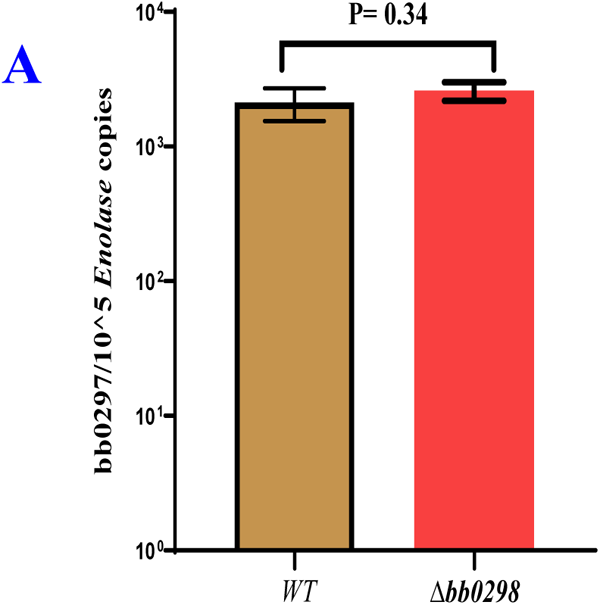
Expression of the gene upstream of the targeted *bb0298* was not affected by the in-frame insertion of Pl-Kan. Measurement of transcripts in the indicated spirochete clones by qRT-PCR. The relative transcript levels of *bb0297* were determined in wild-type and Δ*bb0298* cells and then normalized to the expression of the *B. burgdorferi* enolase gene. The bars represent the mean ± standard deviation of the mean from three independent samples. Statistical analysis was performed by using Student’s *t*-test. P-values between samples are shown at the top. A P-value <0.05 was considered statistically significant.

**Figure S5B.**
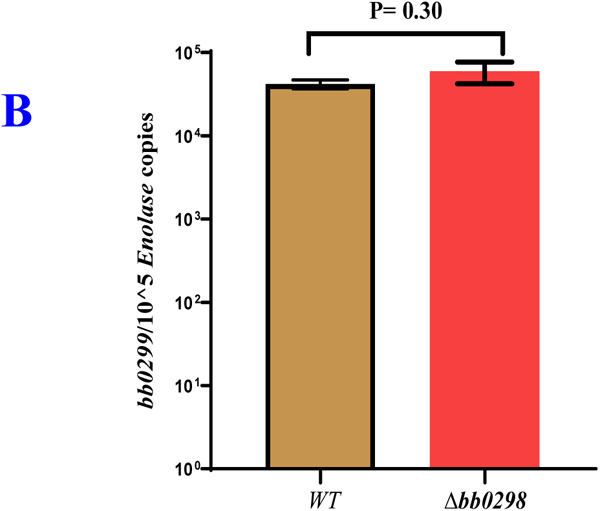
Expression of the gene downstream of the targeted *bb0298* was not affected by the in-frame insertion of Pl-Kan. Measurement of transcripts in the indicated spirochete clones by qRT-PCR. The relative transcript levels of *bb0299* were determined in wild-type and Δ*bb0298* cells and then normalized to the expression of the *B. burgdorferi* enolase gene. The bars represent the mean ± standard deviation of the mean from three independent samples. Statistical analysis was performed by using Student’s *t*-test. P-values between samples are shown at the top. A P-value <0.05 was considered statistically significant.

**Figure S6.**
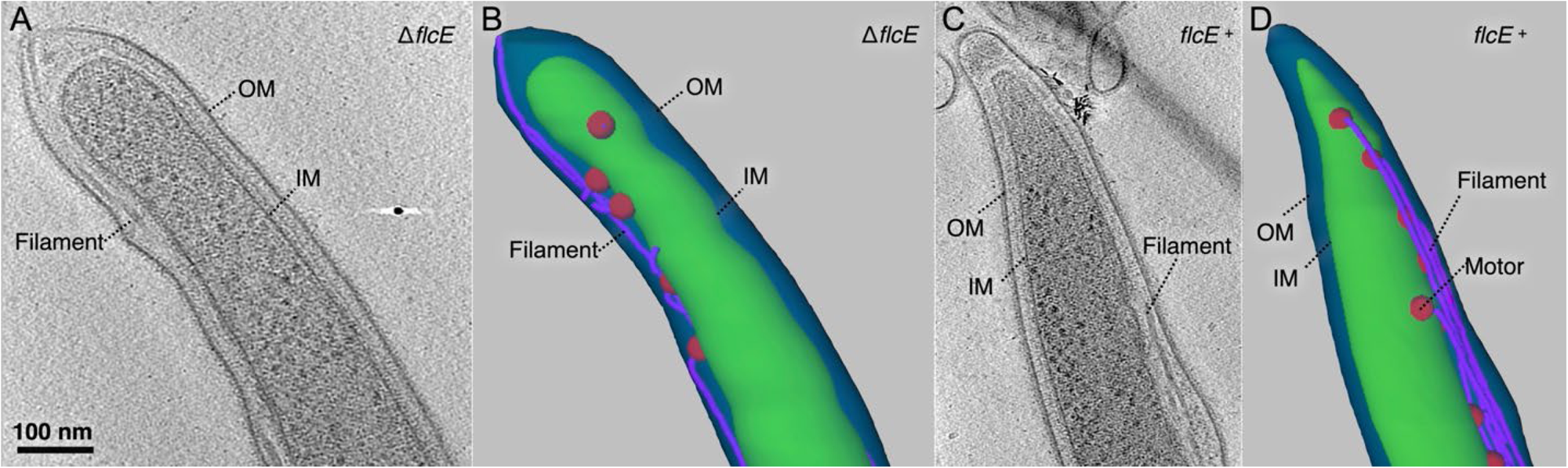
Visualization of Δ*flcE* and *flcE*+ mutants by cryo-ET. **(A)** Slice from a cryo-tomogram showing the tip of a Δ*flcE* cell. The outer membrane (OM), inner membrane (IM), and flagellar filament (Filament) are indicated. **(B)** Corresponding 3D rendering of the Δ*flcE* cell tip in (A), highlighting the flagellar motors (red), filaments (purple), outer membrane (blue), and inner membrane (green). **(C)** Slice from a cryo-tomogram showing the tip of a *flcE*+ cell. The flagellar motor (Motor) is labeled. **(D)** Corresponding 3D rendering of the *flcE*+ cell tip shown in (**C**).

**Figure S7.**
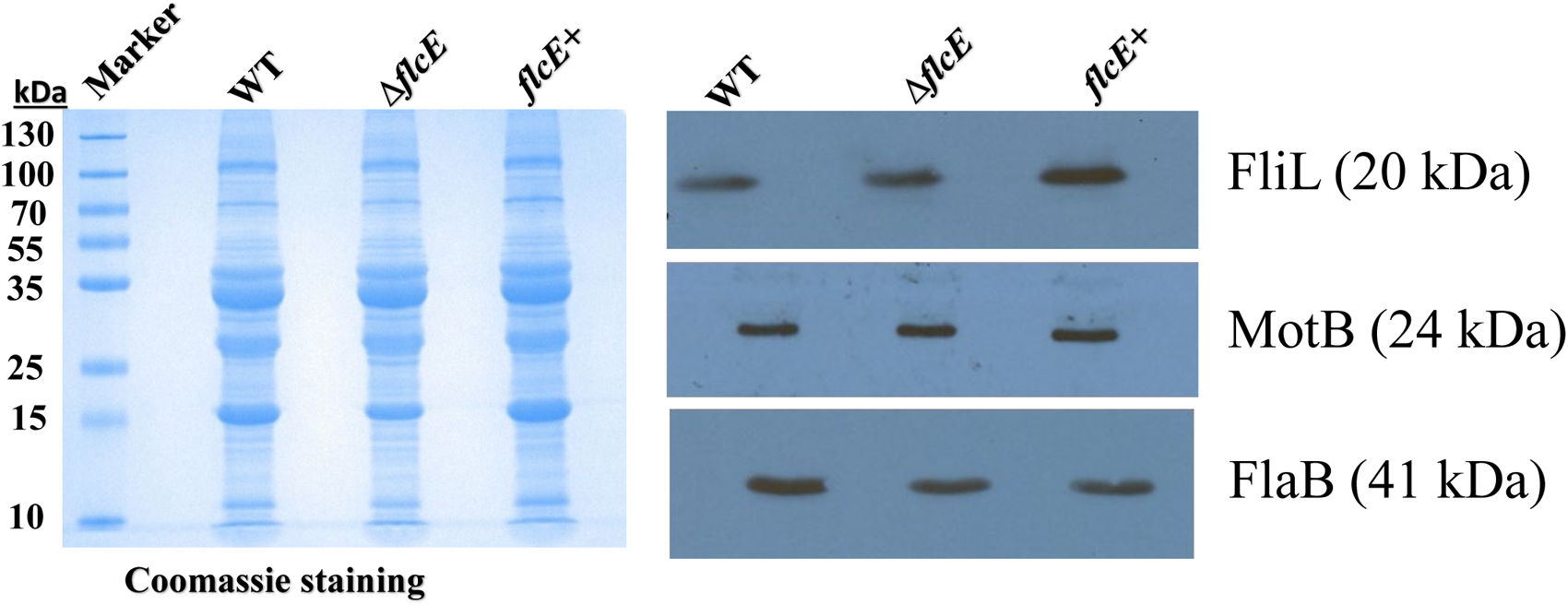
Equal amounts (10 μg) of cell lysates from exponentially growing *Borrelia burgdorferi* were processed for SDS-PAGE, and the gel was then stained with Coomassie blue. A parallel SDS-PAGE gel containing the separated protein lysates was transferred to a PVDF membrane and subsequently immunoblotted separately using antibodies specific to *B. burgdorferi* FliL, MotB, and FlaB. FlaB was utilized as a loading control

**Figure S8.**
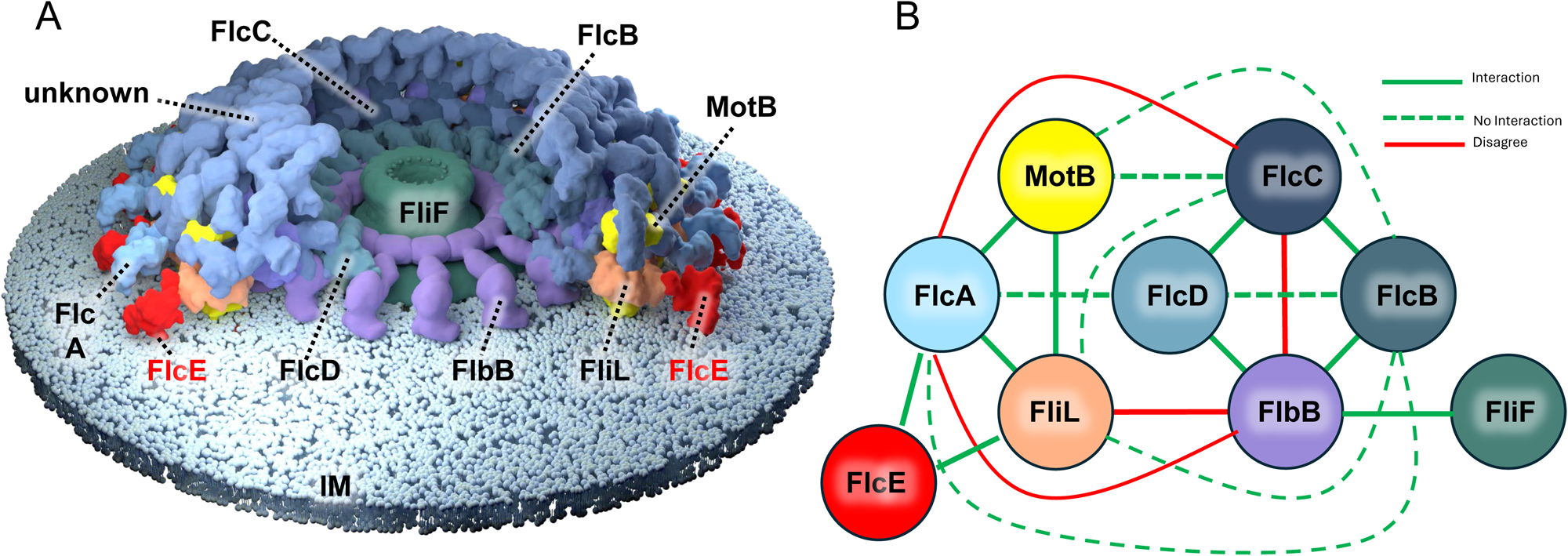
FlcE introduces additional connectivity of the collar-FliL-stator interface. **(A)** Surface-rendered pseudoatomic model depicting the detailed organization of the periplasmic flagellar collar and associated components. Many components of the collar remained unidentified (unknown). **(B)** Experimentally validated protein–protein interaction network of the identified *B. burgdorferi* flagellar collar proteins. Most model-predicted interactions align with the experimental results (green lines), while two experimentally observed interactions are not supported by the model (red lines).

---

**Table S1.**
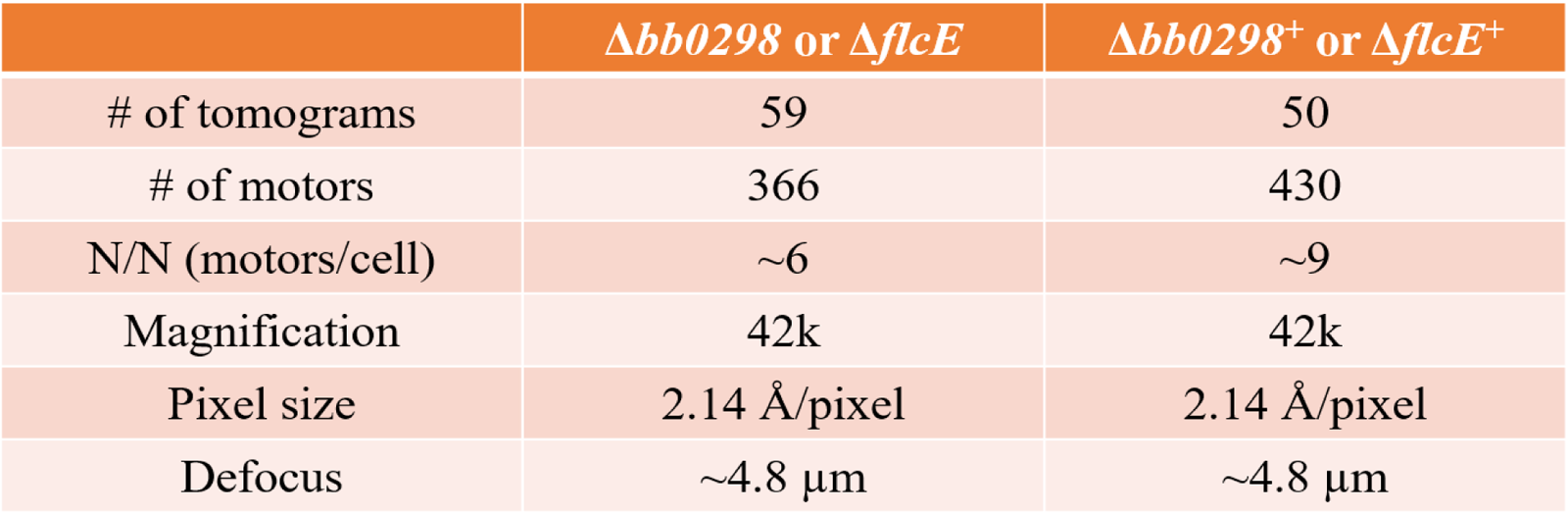
Summary of cryo-ET data collection.

|  | *Δbb0298* or *ΔflcE* | *Δbb0298*<sup>+</sup> or *ΔflcE*<sup>+</sup> |
| --- | --- | --- |
| # of tomograms | 59 | 50 |
| # of motors | 366 | 430 |
| N/N (motors/cell) | ~6 | ~9 |
| Magnification | 42k | 42k |
| Pixel size | 2.14 Å/pixel | 2.14 Å/pixel |
| Defocus | ~4.8 μm | ~4.8 μm |

**Table S2.** Organization of the *dcw* gene cluster in members of Spirochaetales and Winmispirales.

| Organism name | FlcE locus tag,<br>GenBank<br>accession | *dcw* - flagellar gene cluster |  |
| --- | --- | --- | --- |
|  |  | GenBank entry,<br>genomic coordinates | Gene order |
| **Order Spirochaetales** |  |  |  |
| ***Breznakiellaceae* (termite gut spirochetes)** |  |  |  |
| *Breznakiella homolactica*<br>RmG30 | JFL75_10565,<br>QQO07405.1 | CP067089.2:<br>45,617..69,301 | 24 genes: *mraZ-rsmH-ftsL-murF-mraY-ftsW-ftsQ-ftsA-ftsZ-xerD-flcE-dprA-topA-xerC-hslV-hslU-flgB-flgC-fliE-fliF-fliG-fliH-fliI-fliJ* |
| *Gracilinema caldarium*<br>DSM 7334 | Spica_1679,<br>AEJ19822.1 | CP002868.1, reverse:<br>1,870,284..1,893,413 | Same |
| *Leadbetteria azotonutricia*<br>ZAS-9 | TREAZ_RS06765<br>(WP_043922962) | NC_015577.1:<br>1,550,643..1,574,179 | Same |
| ***Spirochaetaceae*** |  |  |  |
| *Oceanispirochaeta*<br>*crateris* K2 | EXM22_07825,<br>QEN07900.1 | CP036150.1, reverse:<br>1,697,967..1,719,290 | 23 genes, no *topA*;<br>after *fliJ*, 18 more flagellar genes |
| *Salinispira pacifica*<br>DSM 27196 | L21SP2_1167,<br>AHC14570.1 | CP006939.1:<br>1,232,468..1,256,047 | 27 genes, no *topA*, inserted *deoC*<br>and three short hypothetical genes:<br>*mraZ-rsmH-ftsL-murF-mraY-ftsW-ftsQ-ftsA-126 nt-ftsZ-xerD-deoC-165 nt-flcE-dprA-xerC-hslV-hslU-flgB-flgC-fliE-fliF-fliG-123 nt-fliH-fliI-fliJ*; then 21 more flagellar genes |
| *Sediminispirochaeta*<br>*smaragdinae* DSM 11293 | Spirs_1554,<br>ADK80681.1 | CP002116.1:<br>1,661,125..1,682,188 | 23 genes, no *topA*;<br>after *fliJ*, 18 more flagellar genes |
| *Spirochaeta africana*<br>DSM 8902 | Spiaf_2113,<br>AFG38161.1 | CP003282.1, reverse:<br>2,386,602..2,407,550 | 24 genes, no *topA*, a hypothetical<br>*195-nt* gene between *xerD* and *flcE*;<br>after *fliJ*, 18 more flagellar genes |
| *Thiospirochaeta perfilievii*<br>DSM 19205 | EW093_03175,<br>QEN03741.1 | CP035807.1, reverse:<br>680,059..700,613 | 23 genes, no *topA*; after *fliJ*, one<br>more flagellar gene, *motE* |
| *Alkalispirochaeta*<br>*sphaeroplastigenens* | AU468_02665,<br>POR04734.1 | LPWH01000007.1,<br>reverse: 15,550..37,364 | 23 genes, no *topA*;<br>after *fliJ*, 16 more flagellar genes |
| *Marispirochaeta*<br>*aestuarii* JC444 | B4O97_13315,<br>ORC34287.1 | MWQY01000014,<br>reverse: 75,168..96,204 | 23 genes, no *topA*;<br>after *fliJ*, 18 more flagellar genes |
| *Entomospira culicis*<br>BR151 | PVA46_05430,<br>WDI36766.1 | CP118181.1:<br>1,148,041..1,159,986 | 11 genes: *rsmH-ftsL-murF-mraY-ftsW-ftsQ-ftsA-ftsZ-xerD-flcE-dprA*;<br>a separate flagellar operon |
| ***Borreliaceae*** |  |  |  |
| *Borrelia burgdorferi*<br>B31 | BB_0298,<br>AAC66650.1 | AE000783.1, reverse;<br>297,038..316,143 | 20 genes, no *mraZ*, *xerD*, *topA*, and<br>*xerC*: *rsmH-ftsL-murF-mraY-ftsW-ftsQ-ftsA-ftsZ-flcE-dprA-hslV-hslU-flgB-flgC-fliE-fliF-fliG-fliH-fliI-fliJ*;<br>after *fliJ*, 18 more flagellar genes |
| *Borrelia hermsii* DAH | BH0298,<br>AAX16815.1 | CP000048.1, reverse;<br>299,727..318,814 | Same |
| **Treponemataceae** |  |  |  |
| *Brucepastera parasyntrophica* RmG11 | K7I13_02265,<br>ULQ60166.1 | CP084606.1, reverse:<br>477,555..499,882 | 23 genes, no *mraY* |
| *Teretinema zuelzeriae* DSM 1903 | K7J14_00820,<br>MCD1653251.1 | JAINWA010000001,<br>reverse:<br>152,231..174,338 | Same |
| *Treponema denticola* ATCC 35405 | TDE_1206,<br>AAS11724.1 | AE017226.1:<br>1,230,092..1,252,475 | Same |
| *Treponema pallidum* subsp. *pallidum* str. Nichols | TP_0392,<br>AAC65376.1 | AE000520.1:<br>408,169..428,859 | 21 genes, no *mraY*, *hslV*, or *hslU*:<br>*mraZ-rsmH-ftsL-murF-ftsW-ftsQ-ftsA-ftsZ-xerD-flcE-dprA-topA-xerC-flgB-flgC-fliE-fliF-fliG-fliH-fliI-fliJ* |
| **Rectinemataceae** |  |  |  |
| *Rectinema subterraneum* | N/A | JBNAWG010000003:<br>reverse, 36,183.. 52,337 | 15 genes: *mraZ-rsmH-ftsL-murF-mraY-ftsW-ftsQ-ftsA-ftsZ-xerD-dprA-topA-xerC-hslV-hslU*; no flagellar genes in the genome |
| **Sphaerochaetaceae** |  |  |  |
| *Parasphaerochaeta coccoides* DSM 17374 | N/A | CP002659.1, reverse:<br>744,520..759,739 | 15 genes: *mraZ-rsmH-ftsL-murF-mraY-ftsW-ftsQ-ftsA-ftsZ-xerD-dprA-xerC-hslV-hslU-flhF*; no flagellar genes in the genome |
| *Sphaerochaeta globosa* str. Buddy | N/A | CP002541.1:<br>1,363,848..1,378,730 | 16 genes: *mraZ-rsmH-ftsL-murF-mraY-ftsW-ftsQ-ftsA-ftsZ-xerD-tauE-dprA-xerC-hslV-hslU-flhF*; no flagellar genes in the genome |
| **Spirochaetales incertae sedis** |  |  |  |
| *Exilispira thermophila* JCM 14728 | JCM14728_12900<br>BGT63363.1 | AP042476.1:<br>a) reverse<br>1,545,381..1,552,577;<br>b) reverse:<br>1,113,153.. 1,123,961 | Two operons: a) 7 genes: *flcE-dprA-topA-xerC-hslU-hslV-DUF364*, and b) 10 genes: *mraZ-rsmH-ftsL-murF-mraY-ftsW-ftsQ-ftsA-ftsZ-ygiQ*; also a separate flagellar operon |
| **Order Winmispirales** |  |  |  |
| **Winmispiraceae** |  |  |  |
| *Rarispira pelagica* 38H-sp 4 | WKV44_08975,<br>MEM5948674.1 | JBCHKQ010000004:<br>141,814..164,764 | 24 genes: *mraZ-rsmH-ftsL-murF-mraY-ftsW-ftsQ-ftsA-ftsZ-xerD-flcE-dprA-topA-xerC-hslV-hslU-flgB-flgC-fliE-fliF-fliG-fliH-fliI-fliJ*; after *fliJ*, 18 more flagellar genes |
| *Winmispira thermophila* DSM 6578 | Spith_0926,<br>AEJ61200.1 | CP002903.1:<br>1,044,808..1,067,874 | Same; after *fliJ*, 17 more flagellar genes |

The respective gene clusters can be retrieved in one step using the URL like https://www.ncbi.nlm.nih.gov/nuccore/CP067089.2?from=45617&to=69301 (GenBank output), or https://www.ncbi.nlm.nih.gov/nuccore/AE000783.1?report=graph&from=297038&to=316143&strand=true (reverse graphical output)

